# White and Brown Adipose Tissue Share a Common Fibro-Adipogenic Progenitor Population

**DOI:** 10.1101/2025.05.28.656577

**Authors:** Hoang Bui, Julia K. Hansen, Valentina Lo Sardo, Andrea Galmozzi

**Affiliations:** Department of Medicine, University of Wisconsin-Madison, School of Medicine and Public Health, Madison, WI; Nutrition and Metabolism Graduate Program, University of Wisconsin-Madison, Madison, WI; Department of Cell and Regenerative Biology, University of Wisconsin-Madison, School of Medicine and Public Health, Madison, WI; Department of Biomolecular Chemistry, University of Wisconsin-Madison School of Medicine and Public Health, Madison, WI; University of Wisconsin Carbone Cancer Center, University of Wisconsin-Madison School of Medicine and Public Health, Madison, WI

## Abstract

Adipose tissue heterogeneity has emerged as a central factor in regulating adipose tissue function in physiology and pathophysiology, yet tools to model and study this diversity *in vitro* remain limited. Here, we performed single-cell RNA sequencing on cultured primary white and brown preadipocytes to assess how *in vitro* conditions impact progenitor identity. We identified two major subpopulations in both depots: committed adipogenic precursors (CAPs) and fibro-adipogenic progenitor-like cells (FAPLs). Remarkably, FAPLs were also present in brown adipose tissue, expanding the known landscape of progenitor populations in this depot. Trajectory and regulon analyses revealed that both white and brown FAPLs exhibit similar pro-fibrotic, stress-responsive signatures and diverge early from proliferating progenitor states. Integration of datasets showed that FAPLs from both depots cluster together, emphasizing their conserved identity, while CAPs remain depot-specific. Comparison to previously published *in vivo* single-cell datasets revealed that these *in vitro* populations, including brown adipose FAPLs, correspond to adipose-resident progenitor subtypes, validating the physiological relevance of this model for studying adipose tissue heterogeneity and development.

## Introduction

Beyond its fundamental role in storing and releasing energy, adipose depots substantially impact systemic physiology by means of endocrine signaling, regulation of inflammatory processes, and behavioral modulation(Chouchani & Kajimura, 2019; Ouchi *et al*, 2011; Stern *et al*, 2016). The adipose tissue is critical for systemic insulin sensitivity and glucose homeostasis, with impairments in adipocyte function tightly linked to the onset of obesity and type 2 diabetes (T2D)(Rosen & Spiegelman, 2006; Samuel *et al*, 2010). The two primary types of adipose tissue, white and brown adipose, exhibit distinct functional characteristics. White adipose tissue primarily stores energy in the form of fat, while brown adipose tissue actively participates in thermogenesis and energy expenditure(Pond, 1992). With new cutting edge methodologies, it is now well-established that resident cell types like mature lipid-storing adipocytes, adipocyte precursor cells (APCs), and immune cells represent a heterogeneous population that collectively contributes to the overall functions and remodeling of fat depots in health and disease(Duerre & Galmozzi, 2022; Wang *et al*, 2022). Recent studies have further delineated the complexity within adipose tissues, identifying multiple subpopulations of adipocytes with distinct functions within both mouse and human adipose tissues. For example, LGAs/AdipoPLIN populations consist of insulin-responsive adipocytes involved in lipid synthesis, while LSAs/AdipoLEP populations rely more on lipid uptake rather than *de novo* synthesis(Backdahl *et al*, 2021; Sarvari *et al*, 2021). In interscapular brown adipose tissue (iBAT), two discrete subsets of brown adipocytes, high (BA-H) and low (BA-L) thermogenic adipocytes, respectively, have been identified based on the uneven expression of adiponectin and the thermogenic protein Ucp1(Cinti *et al*, 2002; Song *et al*, 2020; Spaethling *et al*, 2016). Furthermore, a third class of mature brown adipocytes capable of regulating thermogenesis of neighboring cells by inhibiting thermogenic activity via production and secretion of acetate has been recently discovered in humans and mice(Sun *et al*, 2020; Sun *et al*, 2021).

Akin to fully mature fat cells, adipocyte progenitors (APCs) are a heterogeneous population that plays an integral part in the maintenance of healthy adipose tissue. Many studies applied single-cell RNAseq profiling in mice and humans at different ages and metabolic conditions (i.e., lean vs obese) to study the composition of WAT depots and led to the identification of as many APC subtypes. Fueled by slightly distinct cell sorting strategies, these APC clusters have been tagged with unique names, including adipogenesis-regulatory cells (Aregs)(Schwalie *et al*, 2018), Fibro-inflammatory Progenitors (FIPs)(Hepler *et al*, 2018), Fibro-adipogenic Progenitors (FAPs)(Sarvari *et al*., 2021), or Dpp4^+^ multipotent progenitors(Burl *et al*, 2018; Merrick *et al*, 2019; Rondini *et al*, 2021). Despite this initial lack of consensus, recent efforts have tried to harmonize the nomenclature of white progenitor subtypes, resulting in two main classes consisting of preadipocytes, or Committed Adipocyte Progenitors (CAPs), and Fibro-Adipogenic Progenitors (FAPs)(Maniyadath *et al*, 2023). Conversely, the spectrum of brown adipocyte progenitors is less understood. One study identified several subtypes, primarily localized around the vasculature, including the adipogenic quiescent/cold-responsive ASC1 alongside less-defined ASC2 and ASC3(Burl *et al*., 2018). However, whether these cells represent distinct progenitor subtypes or reflect different stages of maturation of the same brown adipocyte progenitors remains unclear(Karlina *et al*, 2021).

We recently optimized a method to isolate primary white and brown preadipocytes from neonatal mice(Galmozzi *et al*, 2021). Adipose depots of newborn animals are rapidly growing and are enriched in progenitor cells with high proliferative capacity and high differentiation potential(Wang *et al*, 2013). Here, we conducted single cell RNAseq analysis in cultured primary white and brown preadipocytes to determine how accurately these *in vitro* models recapitulate the heterogeneity of adipocyte progenitors *in vivo*. Consistent with previous reports, we identified two main preadipocyte subpopulations for white preadipocytes consisting of committed adipogenic progenitors (wCAPs) and fibro-adipogenic progenitors (FAPs-like, or wFAPLs). Similarly, brown preadipocytes also display a bifurcated differentiation commitment, with one subpopulation of classical adipocyte precursors (bCAPs) and the other one presenting more fibro-inflammatory characteristics (bFAPLs). Notably, we show that wFAPLs and bFAPLs are remarkably similar and exhibit identical pro-fibrotic properties, suggesting a shared regulatory role in BAT and WAT function.

## Results

### Characterization of white and brown adipocyte precursors

Following isolation from male and female C57/BL6 pups, white and brown adipose tissue progenitors were expanded *in vitro* for 5 days, trypsinized into a single-cell suspension and processed for single-cell RNA sequencing using the 10X Genomics platform (**Figure 1A**). Unbiased clustering confirmed an enrichment of 98.28% and 97.08% for adipocyte precursor cells (APCs, *Pdgfra^+^*, *Pdgfrb^+^*, and *Dlk1^+^*) isolated from white and brown adipose depots, respectively (**Figure 1B**-**E**). Non-adipose cells constituted a minor percentage of the total cell population in both WAT and BAT and were identified as myelin-producing glial cell (*Plp1^+^*, 1.07% in WAT and 1.11% in BAT), myeloid cells (*Ptprc^+^*, 0.66% in WAT and 1.27% in BAT) and myocytes (*Myod1^+^,* 0.54% in BAT) (**Figure 1B-E**). Removal of non-adipose cells resulted in 4331 WAT and 5958 BAT progenitors that were used for downstream analysis (**Figure EV1A** and **EV1F**). To minimize the impact of cell cycle phase, which appeared to be a major determinant in defining cell populations (**Figure EV1B**, **EV1C**, **EV1G**, and **EV1H**), we conducted cell cycle regression, re-clustered WAT (**Figure EV1D** and **E**) and BAT progenitors (**Figure EV1I** and **J**), and identified seven distinct clusters for both WAT (**Figure 1F**) and BAT preadipocytes (**Figure 1G**).

**Figure 1.**
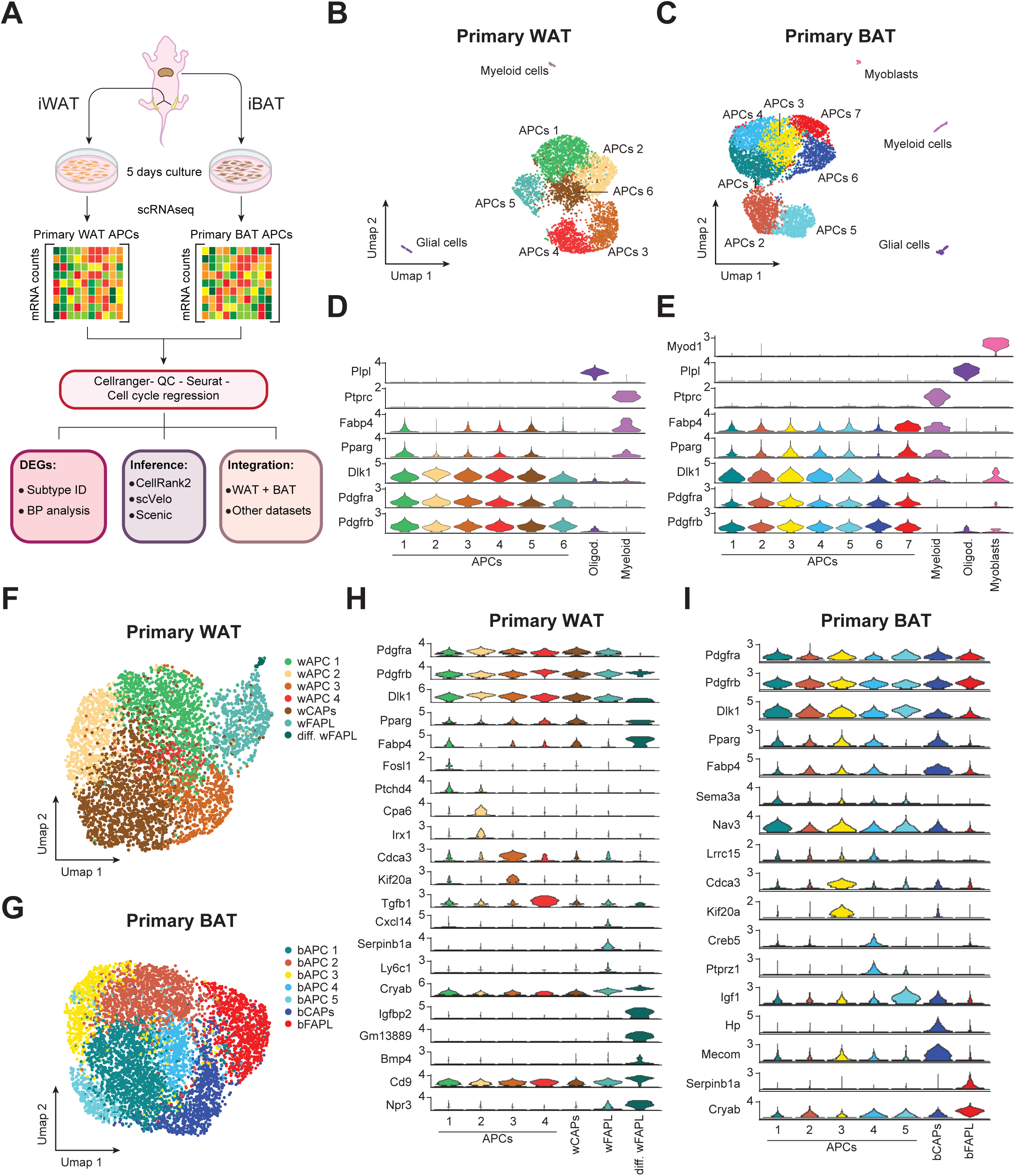
Cultured primary WAT and BAT preadipocytes maintain their heterogeneity. (**A**) Schematic of the single-cell RNA-seq pipeline used to isolate and characterize adipose progenitor cells (APCs) and downstream integrative analyses, including data quality control, cell cycle regression, and subpopulation annotation from neonatal white (WAT) and brown (BAT) adipose tissues. (**B**,**C**) UMAP embeddings showing global clustering of all recovered cell types, including APCs, myeloid cells, glial cells, and myoblasts in WAT (**B**) and BAT (**C**). (**D**,**E**) Expression of representative marker genes (e.g., *Pdgfra*, *Pdgfrb*, *Pparg*, *Dlk1*, *Fabp4*) highlights the identity of non-adipo cells in WAT (**D**) and BAT (**E**). (**F**) UMAPs showing clusters of WAT APCs (n = 4331) after cell cycle regression. (**G**) UMAPs showing clusters of BAT APCs (n = 5958) after cell cycle regression.(**H**) Violin plots of representative markers of wAPC 1-4, wCAPs, wFAPLs, and differentiating wFAPLs. (**I**) Violin plots of representative markers of bAPC 1-5, bCAPs, and bFAPLs.

Like mesenchymal stromal cells(Gao *et al*, 2018; Turley *et al*, 2015; Vishvanath *et al*, 2016), all adipocyte progenitors from both WAT and BAT showed wide expression of platelet-derived growth factor receptors *Pdgfra* and *Pdgfrb* (**Figure 1H** and **I**). Similarly, the common marker of preadipocytes *Dlk1*(Smas & Sul, 1993) was also widely expressed across all WAT and BAT clusters (**Figure 1H** and **I**). Nevertheless, each WAT and BAT cluster was defined by a unique set of differentially expressed genes (**Figure EV2A**, **B** and **Tables EV1** and **EV2**). Specifically, we identified four WAT clusters characterized by low expression of *Pparg* and a distinct gene signature (**wAPC 1**, *Fosl1^+^*and *Ptchd4^+^*, 22.6%; **wAPC 2**, *Cpa6^+^* and *Irx1^+^*, 14.8%; **wAPC 3**, *Cdca3^high^* and *Kif20a*^+^*, 13*%; and **wAPC 4**, *Tgfb1^high^*, 2.9%) (**Figure 1H**). Notably, one cluster showed significantly higher expression of canonical adipocyte markers (i.e., *Pparg^high^*, *Fabp4^high^*) and was therefore labeled as white adipocyte precursors, or **wCAPs** (**Figure 1F** and **H**). Conversely, another cluster, that we called white Fibro-Adipogenic-Progenitor-Like (**wFAPLs**), was characterized by the expression of chemokines (*Cxcl14^+^*), stress response genes (*Cryab^high^*), and inflammation-related genes (*Ly6c1^+^*, *Serpinb1a^+^*), reminiscent of the FAPs previously identified in mouse and human WAT (**Figure 1F** and **H**). Finally, a small percentage of these cells (*Igfbp2^+^*, *Gm13889^+^, Bmp4*^+^, *Cd9^high^*, *Npr3^high^*) also showed high expression of *PPARg^+^* and *Fabp4^+^*, suggesting that they were likely captured during their differentiation into mature adipocytes (**Figure 1F** and **H**). Supporting this hypothesis, this cluster also displayed the lowest expression of *Pdgfra* and *Dlk1* (**Figure 1H**). However, because their overall transcriptional signature closely aligns with wFAPLs, we named this cluster **differentiating wFAPLs** (**Figure 1F** and **H**).

Similarly, each BAT cluster had a unique set of differentially expressed genes (**Figure EV2B** and **Table EV2**). Of them, five BAT clusters (**bAPC 1**, *Sema3a^high^*, *Nav3^high^,* 26.8%; **bAPC 2,** *Lrrc15^+^*, 21.5%; **bAPC 3**, *Cdca3^+^*, *Kif20a^+^,* 9.85%; **bAPC 4**, *Creb5^+^*, *Ptprz1^+^*, 7.42%; and **bAPC 5**, *Igf1^high^*, *6%)* showed low expression of *Pparg* and its target genes (e.g., *Fabp4*) and high *Dlk1* levels, suggesting an early stage of brown adipose tissue progenitors (**Figure 1G** and **I**). As observed for WAT progenitors, one BAT population showed higher expression of *Pparg* and *Fabp4* (*Pparg^high^*, *Fabp4^high^*) and represented a cluster of brown committed adipocyte precursors (**bCAPs**) (**Figure 1G** and **I**). Interestingly, in BAT, a cluster representing ∼15% of the total cell population was characterized by low *Pparg* and *Fabp4* levels and high expression of ECM/stress/immunomodulatory genes (*Serpinb1a^+^*, *Cryab^+^*). Given their similarity to wFAPLs, we named this cluster brown Fibro-Adipogenic-Progenitor-Like cells (**bFAPLs**) (**Figure 1G** and **I**).

### WAT progenitors display two distinct developmental trajectories *in vitro*

To determine whether white progenitor clusters constitute discrete cell types or different transition states of the same progenitors, we performed pseudotime analysis to reconstruct the differentiation trajectories of white preadipocytes using RNA velocity ScVelo(Bergen *et al*, 2020) and define macrostates using CellRank(Lange *et al*, 2022). ScVelo identified two major differentiation paths: one leading to wCAPs through wAPC2, and the other to differentiating wFAPLs via wFAPLs (**Figure EV3A**). Additionally, CellRank identified three macrostates, consisting of wAPC1, wCAPs, and differentiating wFAPLs (**Figure 2A** and **Figure EV3B**). Notably, all three macrostates exhibited metastability scores (i.e., the likelihood to remain in a given state short-term) consistent with terminal state classification (**Figure EV3C**). However, the low stationary distribution, indicating a reduced tendency to remain in that state in the long-term, of wAPC 1 also identified it as the only initial macrostate (**Figure 2A** and **Figure EV3C**). This suggests that the wAPC 1 cluster may serve as a reservoir of proliferating adipocyte progenitors capable of either maintaining an undifferentiated state (i.e., remaining as wACP1) or progressing along the pseudotime trajectory towards wCAPs via the intermediate state wAPC 2 or towards differentiating wFAPLs via their earlier state wFAPLs (**Figure 2B** and **EV3D**).

**Figure 2.**
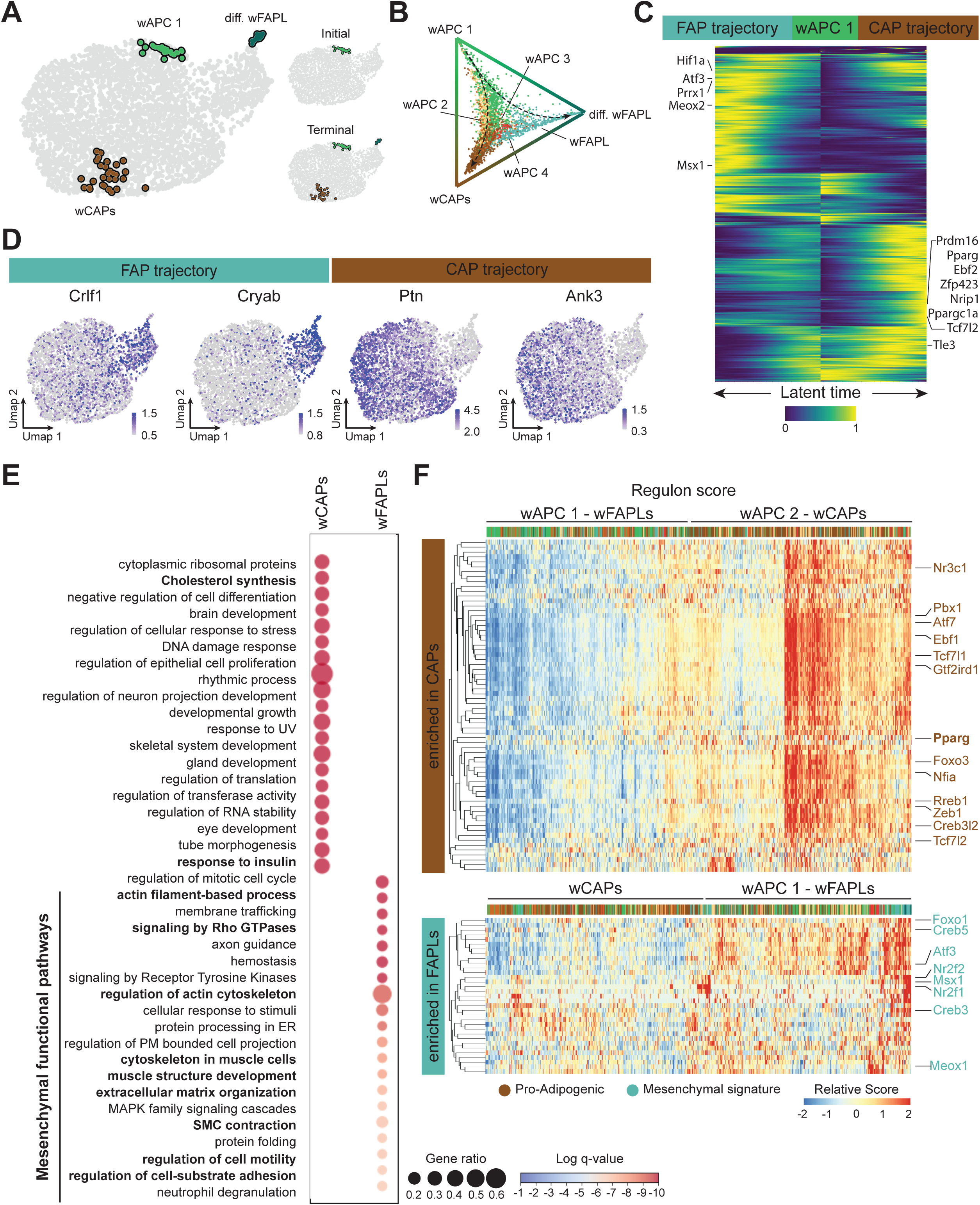
Developmental trajectories of WAT progenitors. (**A**) CellRank analysis reveals 3 macrostates: wAPC1, wCAPs, and differentiating wFAPLs. While all 3 states identified as terminal states (bottom right), only wAPC 1 is also identified as initial state (top right). (**B**) Fate probabilities for each cell towards the three terminal states, highlighting wCAPs and wFAPLs as true distinct terminal lineages. (**C**) Heatmap of the driver genes in wCAP and wFAPL trajectories, computed via generalized additive modeling (GAM). Genes are ordered by their temporal activation patterns along inferred latent time. (**D**) Feature plots of representative drivers overlapping with wCAPs and wFAPLs DEGs, respectively, underscore the divergence between wFAPLs and wCAPs fate. (**E**) Biological pathway analysis of differentially expressed genes between wCAPs vs wFAPLs reveals enrichment in metabolic and insulin responsive pathways in wCAPs, while mesenchymal-related pathways are enriched in wFAPLs. (**F**) Hierarchical heatmaps of regulon scores centering transcription factors enriched in wCAPs (top, defined by wCAP differential scores) and wFAPLs trajectories (bottom, defined by wFAPL differential scores), showing alignment of subclusters to their respective fates.

Having identified the initial and terminal fates, we next sought to visualize gene expression dynamics along these lineages to uncover key regulatory factors. To accomplish this, we computed putative driver genes for each of the two fates (**Table EV3**), plotting them side by side based on their temporal ordering (**Figure 2C**). Interestingly, 227 transcriptional regulators (8.4% of total drivers), including known regulators of adipogenesis such as *Pparg*, *Prdm16*, *Tcf7l2*, *Tle3*, *Zfp423*, *Ebf2*, *Nrip1*, and *Ppargc1a*, were observed amongst the drivers of wCAPs (**Figure 2C**, **Figure EV3E** and **Table EV3**). Conversely, only 112 transcriptional regulators (5.1% of total drivers), mostly linked to mesenchymal cell proliferation and differentiation, were found as drivers of differentiating wFAPLs (**Figure 2C**, **Figure EV3E** and **Table EV3**).

To further characterize cluster-defining genes, we performed biological pathway analysis of genes differentially expressed between wCAPs and wFAPLs (**Fig. 2E** and **Table EV3**). Classical adipocyte markers, including *Pparg* and *Fabp4*, insulin growth factors *Igf1* and *Igf2*, and lipid metabolism-related genes such as *Lpl* and *Cyp7b1* were significantly higher in wCAPs, whereas the stress response protein *Cryab*, the ECM-related genes *Col4a1*, *Col4a2*, and *Serpinb1a*, the anti-adipogenic mesenchyme homeobox *Meox2*(Liu *et al*, 2015), and other inflammatory-related genes, including *Crlf1* and *Cxcl14*, were higher in wFAPLs and/or differentiating wFAPLs, strongly suggesting a mesenchymal transcriptional signature (**Table EV3**). More globally, wCAPs were enriched for metabolic pathways such as cholesterol synthesis and response to insulin, both hallmarks of mature white adipocytes (**Figure 2E**). Conversely, wFAPLs displayed pathways linked to extracellular matrix organization, signaling to Rho GTPases, regulation of the actin cytoskeleton, muscle structure development, smooth muscle cell contraction (**Figure 2E**), processes that have been previously shown to impact expansion of white fat in response to HFD and inflammation(Sun *et al*, 2023) and that confirm the mesenchymal identity of this subpopulation.

Finally, to gain further insights onto the transcriptional networks driving wCAPs and wFAPLs differentiation, we calculated the activity score of transcription factors based on co-expression modules of their putative target genes, also called regulons using SCENIC(Aibar *et al*, 2017). Notably, wCAPs were characterized by high regulon activity of Pparγ as well as several other pro-adipogenic transcription factors (Nr3c1, Ebf1, Tcf7l1, Tcf7l2, Zeb1, Creb3l2) and known modulators of Pparγ transcriptional activity (Pbx1, Atf7, Gtf2ird1, Foxo3, Nfia, Rreb1) (**Figure 2F**). Conversely, wFAPLs exhibited a core set of inflammation-related and mesenchymal fate determination regulons, including Creb3 and Creb5, Foxo1, Atf3, Nr2f1 and 2, Meox1, and Msx1 (**Figures 2F**), strongly supporting the mesenchymal identity of wFAPLs. Furthermore, consistent with the inferred developmental trajectories calculated via RNA velocity (**Figure EV3A**), hierarchical clustering of regulon scores showed segregation of wAPC 1 with wFAPLs, and wAPC 2 with wCAPs (**Figure 2F**), hinting that wAPC 2 is a more committed transition state primed into wCAPs differentiation, while wAPC 1 maintain more FAP-like characteristics. These results support a developmental model for APCs in WAT in which a dormant fibrogenic progenitor population (wAPC 1) can diverge into two distinct developmental trajectories: one leading to committed adipogenic precursors (wCAPs) via intermediate states (e.g., wAPC 1 to wAPC2 to wCAPs) and the other one to fibroadipogenic-like cells (wAPC1 to wFAPLs and differentiating wFAPLs) (**Figure EV3D**).

Altogether, our data indicate that our *in vitro* model can capture cell heterogeneity of adipocyte progenitors concordant to known preadipocytes subtypes previously described and, therefore, this model can be leveraged to investigate cell state transition.

### Primary BAT progenitors mirror the developmental trajectories of their white preadipocyte counterparts

After validating the heterogeneity of WAT preadipocytes, we applied the same computational pipeline to determine the developmental trajectories of the less characterized BAT progenitors. Similar to WAT, we observed two distinct differentiation paths for bCAPs and bFAPLs (**Figure EV4A**). CellRank identified three macrostates representing bCAPS, bFAPLs, and a mixed population of bAPC 1 and bAPC 2 (**Figure 3A**, **B** and **Figure EV4B**), indicating that these two clusters may be sufficiently similar to unify into a single macrostate. As found for wAPC 1, the bAPC 1/2 macrostate was assigned both initial and terminal state (**Figure 3A**, **B**) based on its metastability and stationary distribution (**Figure EV4C**), hinting at bAPC 1 and bAPC 2 as recruitable sources supporting bCAPs and bFAPLs maintenance, while bAPC 3-5 depict intermediate transition states (**Figure 3B** and **EV4D**).

**Figure 3.**
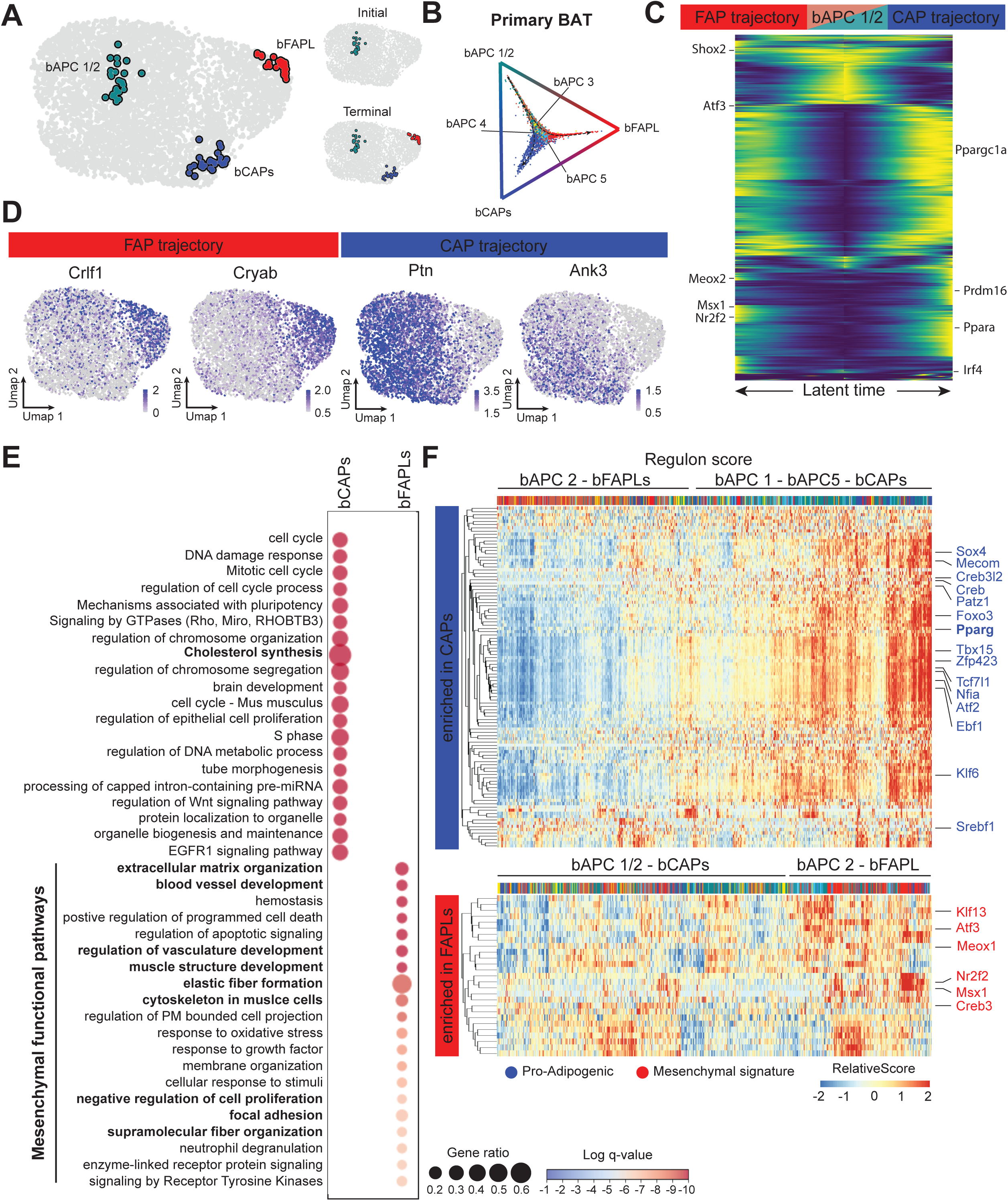
Developmental trajectories of BAT progenitors. (**A**) CellRank analysis reveals 3 macrostates: bAPC1/2, bCAPs, and bFAPLs. While all 3 states identified as terminal states (bottom right), only bAPC 1/2 is also identified as initial state (top right). (**B**) Fate probabilities for each cell towards the three terminal states, highlighting bCAPs and bFAPLs as true distinct terminal lineages. (**C**) Heatmap of the driver genes in bCAP and bFAPL trajectories, computed via GAM. Genes are ordered by their temporal activation patterns along inferred latent time. (**D**) Feature plots of representative drivers overlapping with bCAPs and bFAPLs DEGs, respectively, underscore the divergence between bFAPLs and bCAPs fate. (**E**) Similar to WAT APCs, biological pathway analysis of differentially expressed genes between bCAPs vs bFAPLs shows metabolic pathways in bCAPs, and mesenchymal signature in bFAPLs. (**F**) Hierarchical heatmaps of regulon scores centering transcription factors enriched in bCAPs (top, defined by bCAPs differential scores) and bFAPLs trajectories (bottom, defined by bFAPLs differential scores), showing alignment of subclusters to their respective fates.

Differentially expressed genes and driver gene modules for bCAPs and bFAPLs reflected the differences found in WAT progenitors (**Figure 3C**, **D** and **Table EV4**). The drivers of bCAPs were enriched in pro-adipogenic and thermogenic regulators, such as *Ppara*, *Prdm16*, *Irf4*, and *Ppargc1a* (**Figure 3C**, **Figure EV4E**, and **Table EV4**), whereas drivers of bFAPLs showed mesenchymal and skeletal muscle regulators, including *Msx1*, *Meox1*, *Shox2*, *Nr2f2* and *Atf3* (**Figure 3C**, **Figure EV4E**, and **Table EV4**). Several wFAPLs drivers were also found to be significantly higher in bFAPLs, including *Cryab*, *Meox2*, *Col4a1*, *Col4a2*, *Crlf1*, *Serpinb1a*, and *Nr2f2*, indicating that BAT also possesses a mesenchymal-like progenitor population with fibro-adipogenic characteristics. Consistent with this observation, biological pathway analysis of differentially expressed genes between bCAPs and bFAPLs showed a marked mesenchymal signature of bFAPLs, with ECM remodeling and vasculature development processes (**Figure 3E** and **Table EV4**). On the other hand, in contrast to wCAPs, bCAPs showed only moderate enrichment in metabolic pathways and a stronger signature related to development, cell cycle, and organelle biogenesis and organization (**Figure 3E** and **Table EV4**).

Finally, regulon analysis showed a remarkable similarity with WAT progenitors. Pro-adipogenic transcription factors, including Pparγ, Creb, Tcf7l1, Creb3l2, Atf2, Klf6, Zfp423, Ebf1, Patz1, and Tbx15, and known regulators of Pparγ expression (Sox4, Mecom, Foxo3, Nfia) scored highly in bAPC 1 and bCAPs (**Figure 3F**), whereas inflammation- and mesenchymal-related transcription factors, including Msx1, Meox2, Creb3, Atf3, Nr2f2, and Klf13 were enriched in bAPC 2 and bFAPL transcriptional signatures (**Figures 3F**). Collectively, our data support a developmental model for APCs in BAT in which bAPC 1 and bAPC 2 represent early stages of proliferating brown adipocyte progenitors that can differentiate into committed brown adipogenic precursors (bCAPs) and fibroadipogenic-like cells (bFAPLs), respectively (**Figure EV4D**). Most notably, our results indicate that both WAT and BAT show remarkably similar developmental trajectories and highlight the presence of a previously unappreciated white-like, fibro-adipogenic population in BAT.

### WAT and BAT FAPLs are transcriptionally related to each other

To explore regulatory and developmental parallels between white and brown adipocyte precursors, we combined the two datasets into a unified aggregate. As before, we applied cell cycle regression (**Figure EV5A-F**) and identified nine clusters (**Figure 4A**). Three of these clusters (1–3) were predominantly composed of white progenitors (**Figure 4B** and **Figure EV5G**), four clusters were heavily enriched in brown progenitors (4-7), and two clusters (8 and 9) showed even brown/white cells distribution (40% WAT and 60% BAT in cluster 8, and 59% WAT and 41% BAT in cluster 9, respectively) (**Figure 4B** and **Figure EV5G**). Notably, cluster 1 was composed for the most part (63%) of wCAPs and their earlier progenitors wAPC 2 (22%), while bCAPs (70%) and bAPC 1 (9%) constituted the majority of cluster 7 (**Figure 4B**, **C** and **Figure EV5G**). Of the 2 mixed clusters, cluster 9 only included a small number of cells (145 cells total) coming from wAPC 1 (54%), wAPC 3 (4%), and bAPC 1-4 (2%, 6%, 29%, and 4%, respectively) (**Figure 4B**, **C** and **Figure EV5G**). Interestingly instead, cluster 8 was almost entirely made of white and brown FAPLs (34% and 54%, respectively) (**Figure 4B**, **C** and **Figure EV5G**), further supporting that these fibro-adipogenic-like cells from white and brown adipose depots are remarkably similar. Because cluster 8 was mainly made of FAPLs, its distinct markers included ECM- and stress-related genes such as *Cryab, Meox2, Col4a1, Col4a2, Nr2f2,* and *Serpinb1a*, as seen for white and brown FAPLs independently (**Figure 4D**), as well as low levels of classical adipogenic markers like *Pparg* (**Figure 4D**). Importantly, *Hoxc9* and *Hoxc10*, transcription factor members of the *Hox* family of homeobox genes highly expressed in WAT, but not BAT (Brune *et al*, 2016), were detectable in wFAPLs but not bFAPLs (**Figure 4E**). Conversely, the highly specific marker of interscapular BAT, the transcription factor *Zic1* (Walden *et al*, 2012), was expressed in bFAPLs but not wFAPLs (**Figure 4E**). The detection of depot-specific markers confirmed the absence of cross-contamination between adipocyte populations, addressing concerns associated with the isolation of primary cells, particularly brown preadipocytes, due to the surrounding white adipose tissue. These results ruled out the possibility that bFAPLs originated from co-isolated white adipocyte progenitors and reinforce the conclusion that bFAPLs represents a distinct, *bona fide* progenitor population within brown adipose tissue. In fact, despite the majority (474 genes) of FAPL markers were shared between white and brown FAPLs (**Figure 4F** and **Table EV5**), analyses of genes differentially expressed in white and brown FAPLs also revealed 148 white- and 325 brown-specific markers (**Figure 4F** and **Table EV5**). Conversely, less markers were shared between white and brown CAPs (473 wCAP-specific, 774 bCAP-specific, and 368 shared), which can also explain why FAPLs clustered together while CAPs segregate into distinct clusters when white and brown progenitors are aggregated.

**Figure 4.**
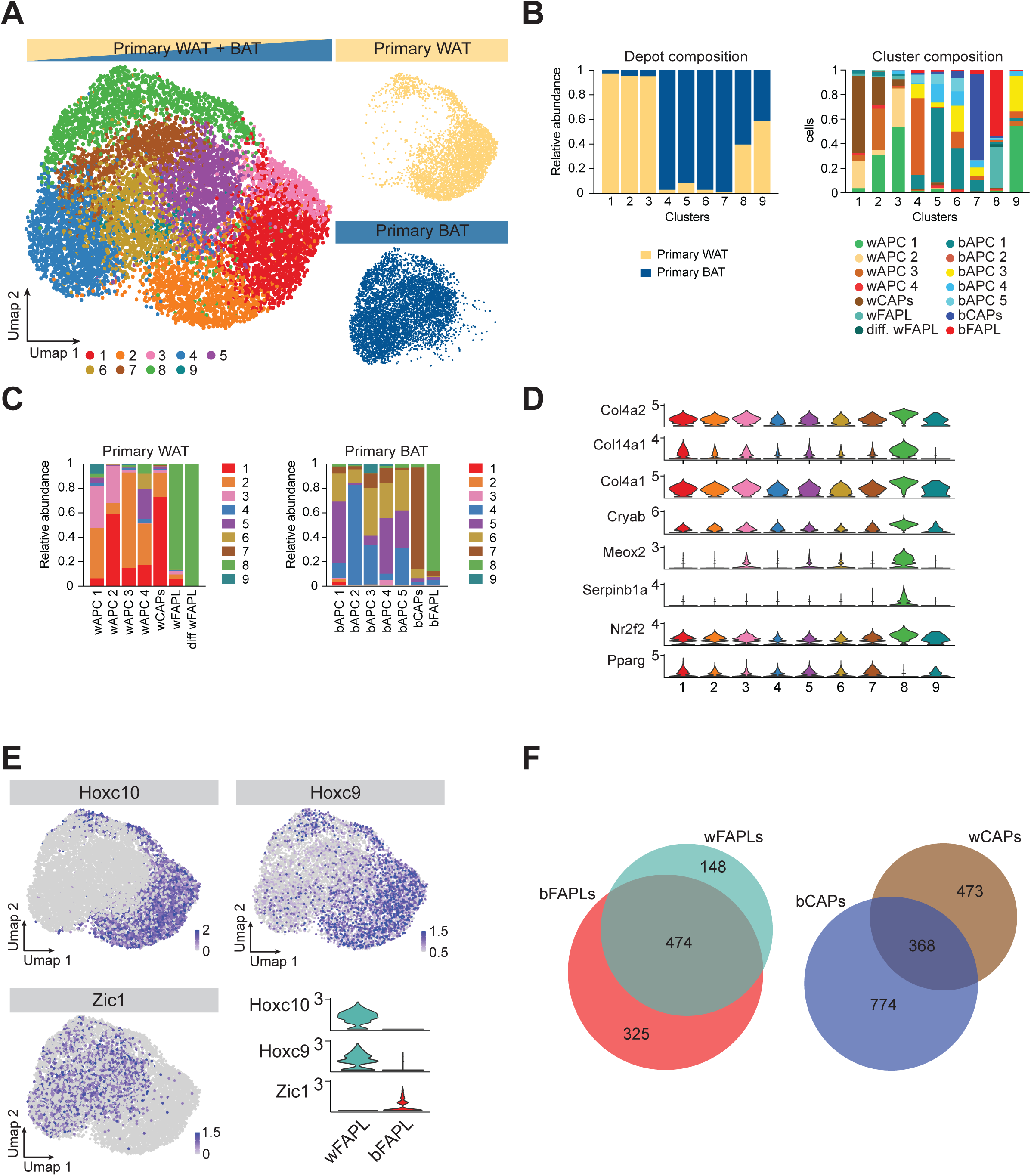
WAT and BAT precursors share a common population of fibro-adipogenic cells. (**A**) UMAP projection of combined WAT and BAT APCs after cell cycle regression reveals nine clusters. Small UMAPs indicate how WAT and BAT cells are distributed. (**B**) Quantification of white and brown APCs across each combined cluster. Clusters 8 and 9 are characterized by equal representation of WAT and BAT cells. (**C**),Quantification of how aggregated clusters compose white and brown APC clusters shows that cluster 8 is made of FAPL progenitors from both WAT and BAT. (**D**) Cluster 8 markers reflect depot-specific FAPL markers. (**E**) Expression of depot-specific genes such as *Hoxc10* and *Hoxc9* for WAT, and *Zic1* for BAT demonstrate absence of cross-contamination of white progenitors in BAT progenitors and vice-versa. (**F**) Venn diagrams representing the overlap of genes differentially expressed between wFAPLs and bFAPLs, and between wCAPs and bCAPs.

Together, these findings indicate that early white and brown progenitors diverge into two distinct lineages: a depot-specific committed adipogenic precursor state (wCAPs/bCAPs), and a shared fibroadipogenic-like state (wFAPLs/bFAPLs) exhibiting a highly conserved transcriptional signature across depots.

### Cultured APCs recapitulate the heterogeneity of APCs *in vivo*

To assess whether our *in vitro* observations align with previously characterized APC populations, we compared our datasets to previously published scRNA-seq/snRNA-seq studies of similar precursor cells(Burl *et al*., 2018; Emont *et al*, 2022; Hepler *et al*., 2018; Holman *et al*, 2024; Karlina *et al*., 2021; Merrick *et al*., 2019; Sarvari *et al*., 2021; Schwalie *et al*., 2018). Somehow surprisingly, some reported markers of adipocyte progenitors, including *Dpp4* and *Icam1*(Merrick *et al*., 2019), *CD142/F3* and *Abcg1*(Schwalie *et al*., 2018), or *Klf4* and *Foxp2* (Sarvari *et al*., 2021), were either expressed at very low levels or not significantly different amongst clusters (**Figure EV6A**). Nevertheless, differentially expressed genes in wFAPLs/differentiating wFAPLs showed substantial qualitative overlap with numerous differentially expressed genes observed in previous studies for mASPC3/mASPC4(Emont *et al*., 2022), FIPs(Hepler *et al*., 2018), FAP1/4(Sarvari *et al*., 2021), G3 (Aregs)/G4 (Schwalie *et al*., 2018), and Dpp4^+^, *Cd142*^+^, or *Spp1*^+^ fibroblasts(Holman *et al*., 2024; Merrick *et al*., 2019) (**Figure EV6B**). Among them, *Fn1*, *Ly6c1*, *Mfap5*, *Creb5*, *Timp2*, *Cd9*, and *Igfbp6* appeared to be consistently elevated in the above-mentioned fibro-adipogenic populations (**Figure EV6B**).

Next, to globally assess the similarity of cultured white and brown progenitors with their counterparts identified *in vivo*, we performed reference mapping using precalculated PCA as integration anchors from our dataset against published datasets(Burl *et al*., 2018; Emont *et al*., 2022; Hepler *et al*., 2018; Holman *et al*., 2024; Karlina *et al*., 2021; Sarvari *et al*., 2021; Schwalie *et al*., 2018). Notably, our wFAPL population shared high qualitative overlap with previously described FAP or FAP-like cells, namely mASPC3 (*Mgp*^+^) and mASPC4 (*Epha3^+^*)(Emont *et al*., 2022), FIPs(*Ly6c1^+^*) (Hepler *et al*., 2018), FAP4 (*Klf4*^+^) and FAP1(*Foxp2*^+^)(Sarvari *et al*., 2021), G3/Aregs (Cd142^+^) and G4 (Ly6c1^+^)(Schwalie *et al*., 2018), and Cd142^+^ fibroblasts(Holman *et al*., 2024) (**Figure 5A** and **B**), indicating that cultured wFAPLs recapitulate the mesenchymal, fibroadipogenic identity observed *in vivo*. Similarly, highly adipogenic mASPC1, mASPC5, and mASPC6(Emont *et al*., 2022), APCs(Hepler *et al*., 2018), FAP2 and 3(Sarvari *et al*., 2021), G1 and G2(Schwalie *et al*., 2018), and *Icam1*^+^ preadipocytes(Holman *et al*., 2024) mapped to wCAPs or their precursors wAPC 2 (**Figure 5A** and **B**), confirming the pro-adipogenic nature of cultured wCAPs.

**Figure 5.**
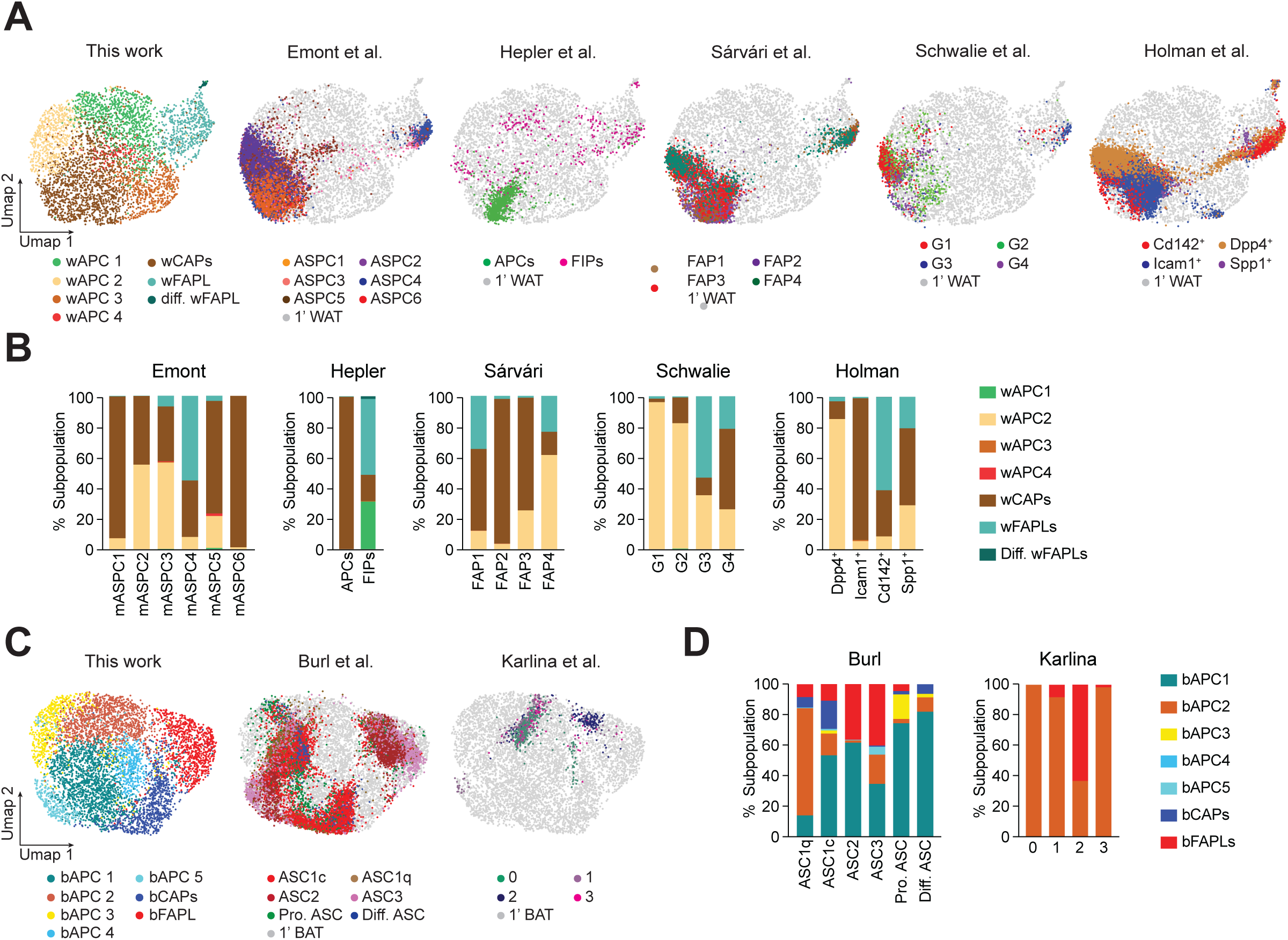
Cultured APCs recapitulate the heterogeneity of APCs found *in vivo*. (**A**) Reference mapping of previously published datasets against primary WAT progenitors links known mesenchymal populations to wFAPLs while highly adipogenic subpopulations map to wCAPs. (**B**) Quantification of reference mapping distribution for WAT datasets. (**C**) Reference mapping of previously published datasets against primary BAT progenitors reveals a fibrogenic population of brown adipose tissue progenitors in *vivo*. (**D**) Quantification of reference mapping for BAT datasets.

Fewer datasets have been generated for BAT progenitors, which have remained to date less thoroughly characterized. Despite this, we found that bCAPs strongly aligned with the previously reported cold-responsive ASC1 progenitors(Burl *et al*., 2018), while bFAPLs overlap in part with the vasculature-associated ASC2/ASC3(Burl *et al*., 2018) (**Figure 5C** and **D**). Similarly, the uncharacterized cluster 2 from another, independent study (Karlina *et al*., 2021) showed significant enrichment (∼60%) of bFAPL-like cells, while all the other clusters (0, 1, and 3) mapped onto bAPC 2, the earlier progenitor of bFAPLs. These results demonstrate that cultured WAT and BAT APCs align with previous *in vivo* phenotypes and that the heterogeneous APC populations isolated from neonatal pups provide a valuable model for investigating distinct adipocyte lineages.

## Discussion

Understanding the cellular heterogeneity of adipose tissue progenitors is central to dissecting the mechanisms regulating adipose tissue development, remodeling, and function. While single-cell RNA-sequencing has recently advanced our knowledge of progenitor populations *in vivo*(Duerre & Galmozzi, 2022; Loft *et al*, 2025; Maniyadath *et al*., 2023), *in vitro* systems that faithfully recapitulate this heterogeneity have remained limited. In this study, we generated single-cell transcriptomic profiles from cultured progenitors derived from both white and brown adipose tissue depots. Our data reveal that key features of adipose progenitor diversity are preserved *in vitro*, providing a powerful tool to bridge *in vivo* findings with mechanistically tractable systems.

A central finding of our work is the resolution of two main progenitor trajectories that are maintained during early *in vitro* culture: committed adipogenic precursors (CAPs) and fibro-adipogenic progenitor-like cells (FAPLs). These populations are transcriptionally distinct and resemble the bifurcating lineages that have been consistently described *in vivo*(Duerre & Galmozzi, 2022; Maniyadath et al., 2023). Importantly, we demonstrate that both white and brown adipose tissue give rise to these two progenitor trajectories, indicating that the underlying lineage programs are conserved across depots. The presence of both adipogenic and fibro-adipogenic lineages under standard culture conditions suggests that *in vitro* models retain a level of progenitor plasticity and heterogeneity relevant for studying adipose tissue physiology.

Most notably, our study reveals a previously undescribed progenitor population in BAT with strong transcriptional similarity to fibro-adipogenic, mesenchymal progenitors found in WAT. FAPs have been well characterized in WAT(Hepler *et al*., 2018; Merrick *et al*., 2019; Sarvari *et al*., 2021; Schwalie *et al*., 2018) and skeletal muscle(Joe *et al*, 2010; Uezumi *et al*, 2010; Wosczyna *et al*, 2019; Yang *et al*, 2022), where they contribute to extracellular matrix production and fibrosis, but had not been described in BAT prior to this study. Using our *in vitro* model and reference mapping to *in vivo* datasets(Burl *et al*., 2018; Karlina *et al*., 2021), we showed that brown adipose FAPs are remarkably similar to white adipose FAPs and are present within brown adipose tissue *in vivo*, highlighting the utility of our system in uncovering progenitor types that may be rare, transient, or previously overlooked in complex *in vivo* environments.

Our dataset also allows alignment of *in vitro*-derived populations with known *in vivo* progenitor classes, including CAPs, Dpp4^+^ interstitial progenitors, FIPs, FAPs, and Aregs. Although APCs dynamically change with aging and pathophysiology(Rondini *et al*., 2021; Sarvari *et al*., 2021; Sun *et al*., 2020; Yang *et al*., 2022; Zhang *et al*, 2022), and direct equivalence between *in vitro* and *in vivo* populations must be interpreted with caution, the consistency in gene expression patterns that we observed between cultured APCs and their *in vivo* counterparts supports the relevance of this *in vitro* system for modeling adipose tissue dynamics. In this context, our results validate a practical and scalable model for dissecting the regulatory mechanisms that govern adipose progenitor fate. Unlike immortalized cell lines, primary cells capture the heterogeneity of APCs *in vivo* and can be confidently used to study how environmental cues or signaling pathways influence adipogenic versus fibro-adipogenic fate. In addition, supporting current efforts to define a broadly accepted nomenclature for adipocyte precursors(Loft *et al*., 2025; Maniyadath *et al*., 2023), this resource might facilitate the identification of shared markers, which may be used for isolation and targeted functional studies, facilitating translational insights into depot-specific remodeling and metabolic disease.

In summary, our study establishes an *in vitro* single-cell atlas of white and brown adipose progenitors that recapitulates key features of *in vivo* biology. By enabling the identification of brown FAPs and providing a framework to study progenitor heterogeneity in controlled conditions, this work offers a valuable resource for the adipose biology community and a platform for mechanistic exploration of lineage dynamics.

## Methods

### Isolation and culturing of primary WAT and BAT progenitors

Primary WAT and BAT progenitors were isolated as previously described(Galmozzi *et al*., 2021). Briefly, subcutaneous white and interscapular brown depots were dissected from male and female C57/BL6 P0 mice into 250 μL ice-cold PBS and 200 μL 2X isolation buffer (123 mM NaCl, 5 mM KCl, 1.3 mM CaCl2, 5 mM glucose, 100 mM 4-(2-hydroxyethyl)-1-piperazineethanesulfonic acid (HEPES), penicillin-streptomycin, and 4% fatty acid-free bovine serum albumin). Tissues were minced using small scissors, then 50 μL of 15 mg/mL collagenase type I was added and incubated at 37°C on a shaker for 35 (WAT) and 50 (BAT) minutes, respectively. After digestion, samples were filtered through a 100 μm cell strainer into 10 mL isolation medium (Dulbecco’s modified Eagle medium (DMEM) + 20% fetal bovine serum (FBS), 10 mM HEPES, 1% penicillin-streptomycin) into a 10 cm dish and incubated at 37°C, 5% CO_2_ for 1.5 hours, followed by three washes with serum-free DMEM to remove blood cells and tissue debris. After washing, 10 mL of isolation medium was added, and cells were incubated overnight. The following day, cells were washed again with serum-free DMEM and maintained in isolation media, which was thereafter refreshed every 2 days. After reaching confluency (in 4 days), cells were washed with PBS and lifted with Trypsin-EDTA for 3 minutes. WAT and BAT progenitor suspensions were then washed in isolation media, counted and live cells processed for single-cell sequencing.

### Single-cell RNA library preparation and sequencing

Single-cell RNA sequencing was conducted by the University of Wisconsin-Madison Biotechnology Center’s Gene Expression Center Core Facility using the 10x Genomics Chromium Single Cell 3ʹ v3.1 platform. Single-cell suspensions were loaded onto the Chromium Next GEM Chip, where cells were partitioned into Gel Bead-In-EMulsions (GEMs) for reverse transcription (RT). Post GEM-RT, cDNA was recovered and purified using DynaBeads MyOne Silane beads (Lot# 160221), followed by amplification through 12 cycles of PCR. The amplified cDNA underwent cleanup with SPRIselect beads (0.6X, Lot# 18198500) and quality was checked using an Agilent HS DNA chip. For library construction, cDNA was fragmented, end-repaired, and A-tailed, with double-sided size selection using SPRIselect beads (0.6x and 0.8x, Lot# 18198500). Adapter ligation was performed, followed by post-ligation cleanup with SPRIselect beads (0.8x). Index PCR was carried out with 7 cycles using the Chromium Dual Index TT Set A (Lot# 160185). The final library was cleaned using SPRIselect beads (0.6x and 0.8x), recovering 35 μL in EB buffer. Library concentration was determined using a Qubit Fluorometer with Qubit dsDNA HS reagents. Final quality control was performed using an Agilent TapeStation with D1000 ScreenTape. Sequencing was initially conducted on a MiSeq Nano, followed by final sequencing on an SP flow cell.

### ScRNAseq Data Analysis

Single-cell RNA-seq data were analyzed by the UW Bioinformatics Resource Center and 10X Cloud services. Quality control of the MiSeq balancing run was performed using UMI-tools. Libraries were balanced for estimated reads per cell and sequenced on an Illumina NovaSeq system. Cell Ranger 7.0.1 software was utilized for demultiplexing, alignment, filtering, barcode counting, UMI counting, and gene expression estimation, using default parameters and the mouse mm10 2020-A genome. High-quality reads were obtained with over 93% mapping confidence. For the WAT sample, 5261 cells were captured with a mean of 48,966 reads per cell and a median of 5777 genes per cell. For the BAT sample, 7009 cells were captured with a mean of 48,249 reads per cell and a median of 5720 genes per cell.

Seurat v5(Hao *et al*, 2024) was used to perform downstream filtering, log-normalization and analyses in R Studio. After applying filters for both WAT and BAT (3900>nFeature RNA> 7700, 1.7%< mitochondrial DNA <5%, 6000< nCount RNA <60000) to remove low quality reads, 4407 cells, 20999 genes for WAT and 6173 cells, 21889 genes for BAT were recovered. Next, non-adipocells were removed from BAT and WAT, resulting in 4331 white precursors and 5958 brown precursors. Cell cycle scoring and regression was applied using default workflow parameters. Cluster analysis at 0.5 resolution with 10 dimensions calculated from the cell-cycle regressed PCA revealed the final clustering, and markers were identified using Seurat FindMarkers function. Biological pathway analysis was performed with Metascape(Zhou *et al*, 2019) while molecular function was assigned using PANTHER classification system (version 19.0).

Public datasets for WAT(Emont *et al*., 2022; Hepler *et al*., 2018; Holman *et al*., 2024; Sarvari *et al*., 2021; Schwalie *et al*., 2018) and BAT(Burl *et al*., 2018; Karlina *et al*., 2021) adipocyte precursors were downloaded and re-processed using Seurat v5. Extraction of data was limited to preadipocytes control on chow diet only. Clustering adjustments were made for the Karlina et al., 2021 and Hepler et al., 2018 datasets to account for cross-platform differences and the availability of clustering parameters. In Hepler et al. dataset, due to the lack of available clustering parameters, the distinction between adipogenic cells (1A and 1B) was consolidated into a single unified APCs group. A clear distinction between APCs and FIPs was observed based on Ly6c1 expression, a key marker for FIPs(Hepler *et al*., 2018). Similarly, the Karlina dataset was reanalyzed using Seurat v5, resulting in highly consistent UMAP projections and clusters.

For reference mapping of public datasets for WAT and BAT against our datasets, common anchors were identified using the FindTransferAnchors function. This function aligns datasets by identifying shared cell features, facilitating integration of disparate datasets based on precomputed PCA space. Next, MapQuery function was used to project the public datasets (query cells) onto our datasets’ UMAPs (reference cells), where 1246 anchors for Emont et al., 373 anchors for Hepler et al., 836 anchors for Sarvari et al., 442 anchors for Schwalie et al., 670 anchors for Holman et al., 1199 anchors for Burl et al., and 265 anchors for Karlina et al. were identified, respectively. The query cells, when mapped, were assigned a predicted identity based on their similarity to the reference clusters, alongside their original labels. The predicted labels were then quantified and represented as subcluster percentages. Finally, to illustrate the spatial alignment, the UMAP coordinates of WAT and BAT datasets with the mapped UMAP coordinates from the published datasets were merged.

For WAT and BAT aggregate, WAT and BAT datasets were combined using merge and reduce function. After combining the two datasets, the cell cycle effect was regressed out of PCA.

### Pseudotime Trajectory Analysis

RNA velocity(Bergen *et al*., 2020) was constructed using the velocyto package, specific to the 10X platform, which generated an RNA velocity (.loom) file for both white and brown adipose tissue. Next, the UMAP coordinates and cluster identities from R/Seurat, together with the matrix reads and the newly generated .loom file, were used as inputs for ScVelo. Differentiation trajectories were analyzed and visualized using the top 2,000 algorithmically defined driver genes using ScVelo’s dynamical modeling alongside Numpy, pandas, scanpy, anndata, igraph, loompy, and matplotlib, following standard workflow. Fate probabilities were assessed via the CellRank2 workflow using the RNA velocity kernel(Lange *et al*., 2022) with time information extracted from ScVelo, and GPCCA(Reuter *et al*, 2019) (pyGPCCA package) for CellRank2 estimators input. The number of macrostates was chosen based on their robustness to parameter changes, and compute_lineage_drivers identified the driver genes of each lineage. To visualize the temporal bifurcation, the significant driver genes (filtered by log-q val<0.05) for each lineage were extracted and their queried gene-expression models at each time point from cr.pl.heatmap were retrieved. Data were compiled into DataFrames and sorted in reverse order for one lineage. Temporal bifurcated expression data were visualized with pheatmap.

### Regulon scoring

Regulon activity was assessed using pySCENIC(Aibar *et al*., 2017), run within a Docker container for reproducibility. The SCENIC+ motif collection(Bravo Gonzalez-Blas *et al*, 2023) was obtained from resources.aertslab.org, alongside mm10 genomic region files centered on 500 bp and 10,000 bp around transcription start sites. Following the SCENIC protocol, pySCENIC first inferred the underlying gene regulatory network, then identified candidate regulons via RcisTarget motif enrichment analysis, and finally scored the activity of these regulons in each cell using the AUCell module. The resulting AUCell score matrix was then merged with the Seurat object for downstream analyses. Subsequent analyses were performed in Seurat and visualized with pheatmap, focusing on transcription factors whose activity patterns appeared especially robust or distinct across clusters through hierarchical clustering.

## Acknowledgements

We thank Dr. Caroline Alexander, Dr. Huy Q. Dinh, Dr. Dave Harris, and current members of the Galmozzi lab for critical input and discussion; the Gene Expression Center Core Facility at the University of Wisconsin-Madison Biotechnology Center for assistance with scRNA sequencing. This work was supported by the National Institute of Health grants 1R35GM150899 (AG) and the Department of Medicine Pilot Funding Program (AG) at the University of Wisconsin-Madison School of Medicine and Public Health. HB is supported by NIH NCATS awards to UW-ICTR TL1TR002375 and UL1TR002373. The model in Figures 2 and 3 were created using BioRender.

## Author Contribution

AG conceived the project. HB, VLS, and AG designed the analyses. HB performed scRNAseq data analysis. JKH performed isolation of primary WAT and BAT progenitors. HB and AG wrote the manuscript and integrated comments from the other authors.

## Data Availability

All data generated via scRNA-seq will be publicly available upon publication and will be deposited in the Gene Expression Omnibus database under Gene Expression Omnibus accession number XXX.

---

**Figure EV1.**
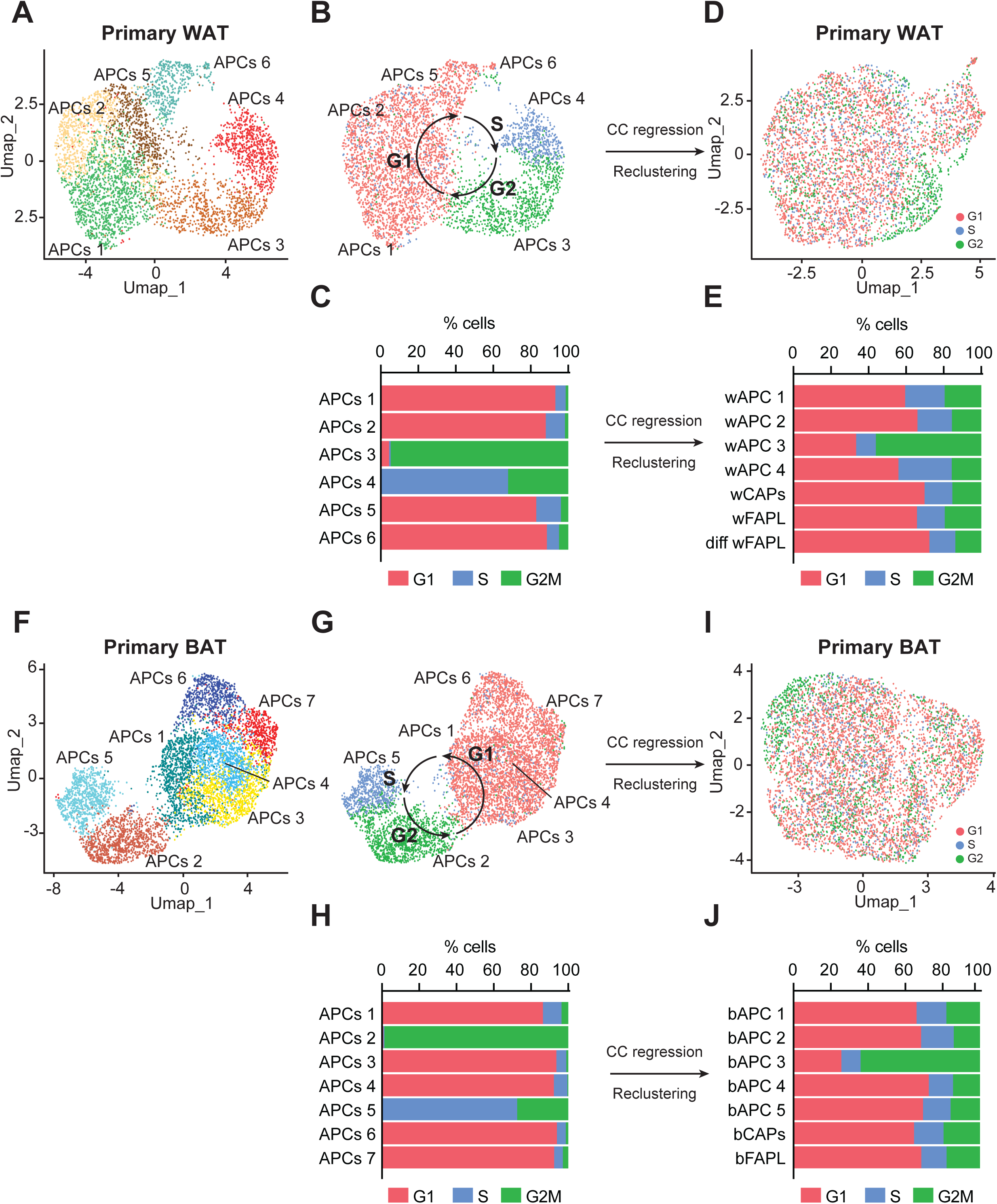
Cell cycle regression and re-clustering of WAT and BAT APCs. (**A**) Clustering of WAT APCs before cell cycle regression. (**B**) Cell cycle phase of WAT APCs. (**C**) Distribution of cell cycle in WAT APC clusters from (**A**). (**D**) Cell cycle regression leads to redistribution of WAT APCs. (**E**) Cell cycle phases are equally distributed across clusters upon cell cycle regression. (**F**) Clustering of BAT APCs before cell cycle regression. (**G**) Cell cycle phase of BAT APCs. (**H**) Distribution of cell cycle in BAT APC clusters from (**F**). (**I**) Cell cycle regression leads to redistribution of BAT APCs. (**J**) Cell cycle phases are equally distributed across BAT clusters upon cell cycle regression.

**Figure EV2.**
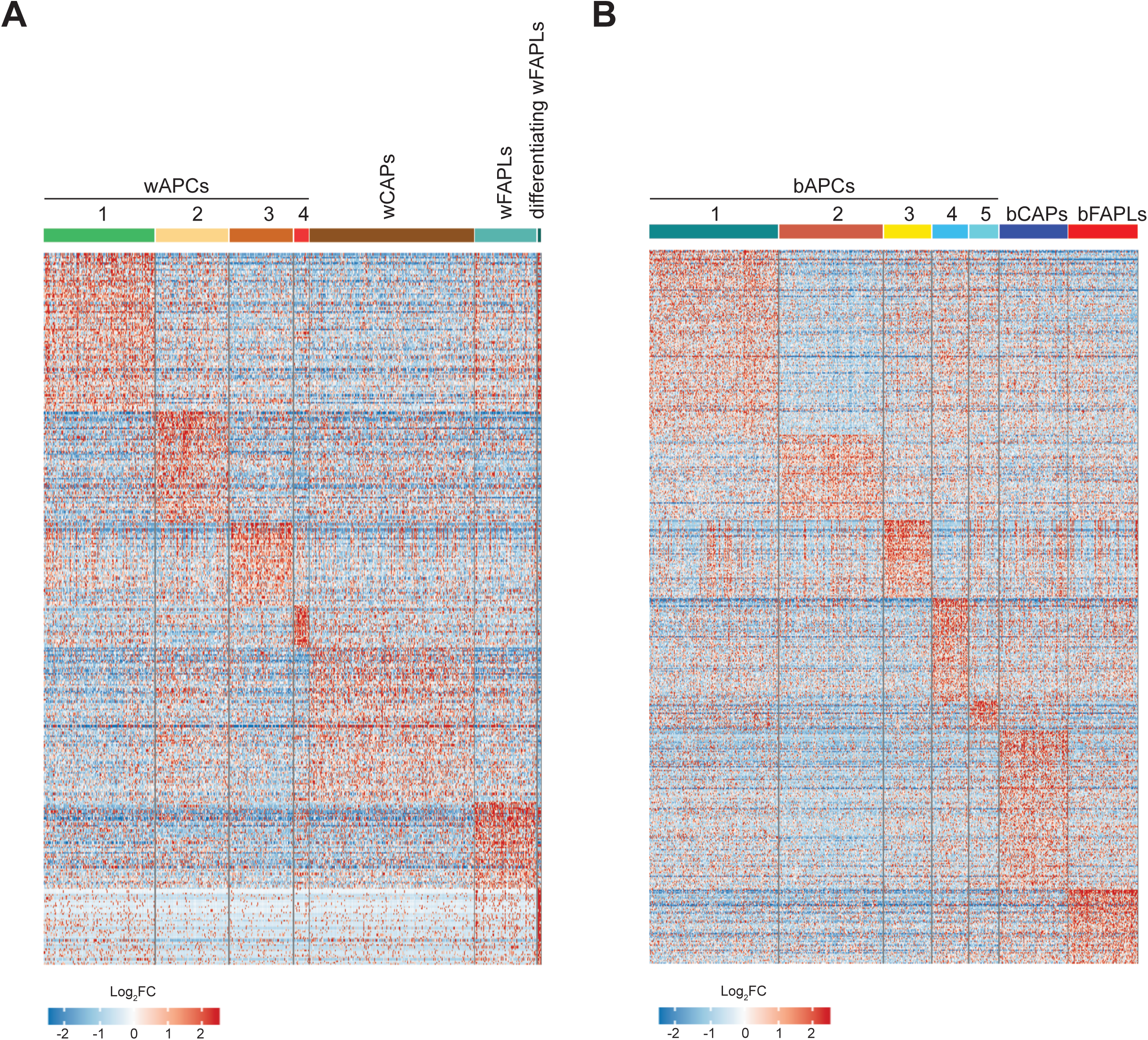
WAT and BAT APC display unique transcriptional signatures. (**A**) Heatmap of the top 10% differentially expressed genes for each white APC cluster. (**B**) Heatmap of the top 10% differentially expressed genes for each brown APC cluster.

**Figure EV3.**
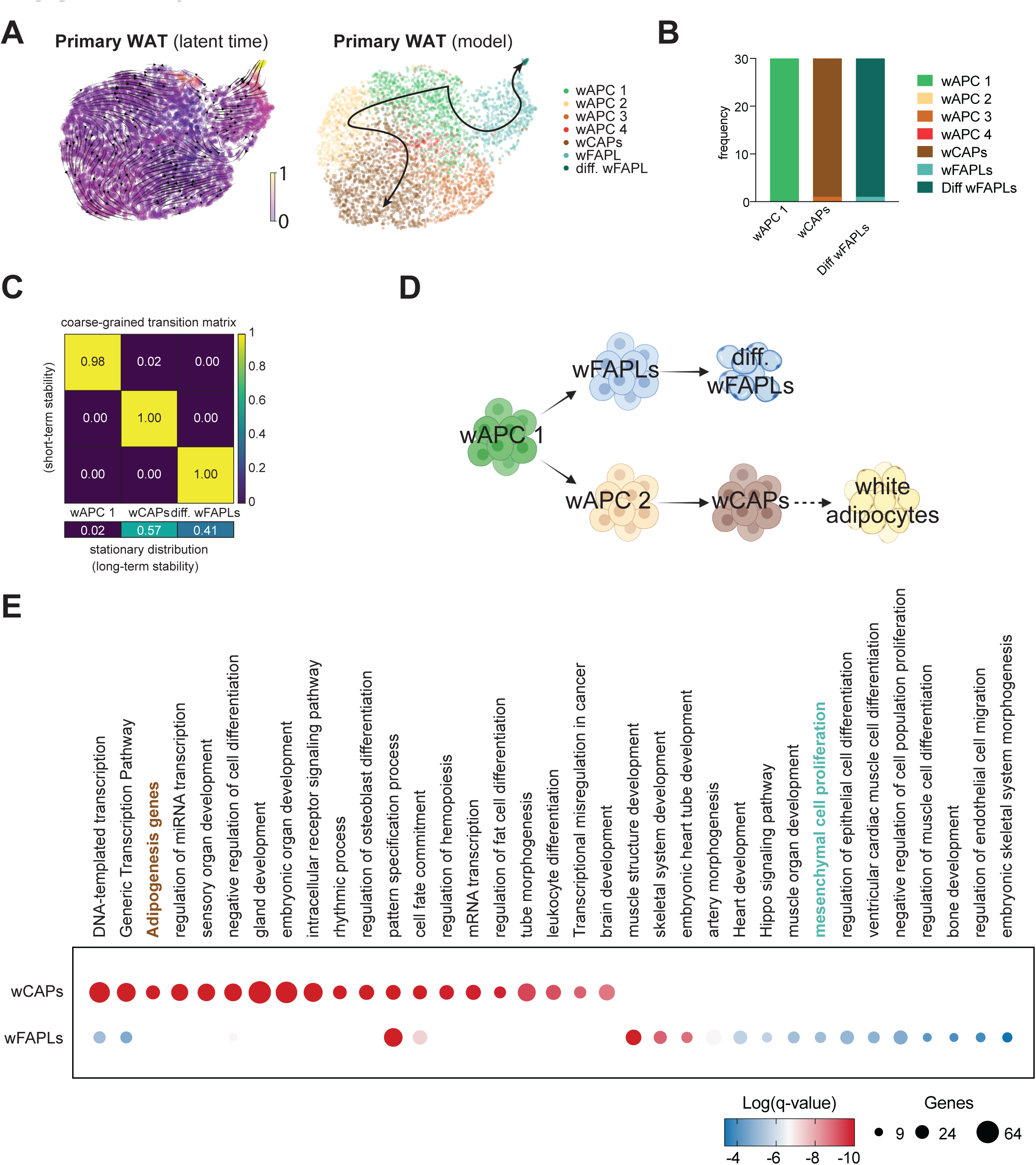
Pseudotime trajectory inferences and macrostate assignments in WAT. (**A**) Latent time UMAP visualization inferred by ScVelo and model of WAT APC differentiation trajectories highlight the transition from early precursors (wAPC 1) to wCAPs and wFAPLs. (**B**) Composition of WAT APC macrostates (n = 30 cells) by CellRank2. (**C**) Coarse-grained transition matrix using CellRank2 GPCCA for WAT macrostates indicating the stability (diagonal elements) and transition probabilities (stationary distribution) amongst macrostates. (**D**) Proposed developmental model of WAT APCs shows a shared early progenitor wAPC 1 that can differentiate into wFAPLs or wCAPs via intermediate transition states (wAPC 2). (**E**) Biological pathway analysis of the transcription factor drivers of wCAPs and wFAPLs shows enrichment in adipogenesis for wCAPs and mesenchymal-related pathways for wFAPLs.

**Figure EV4.**
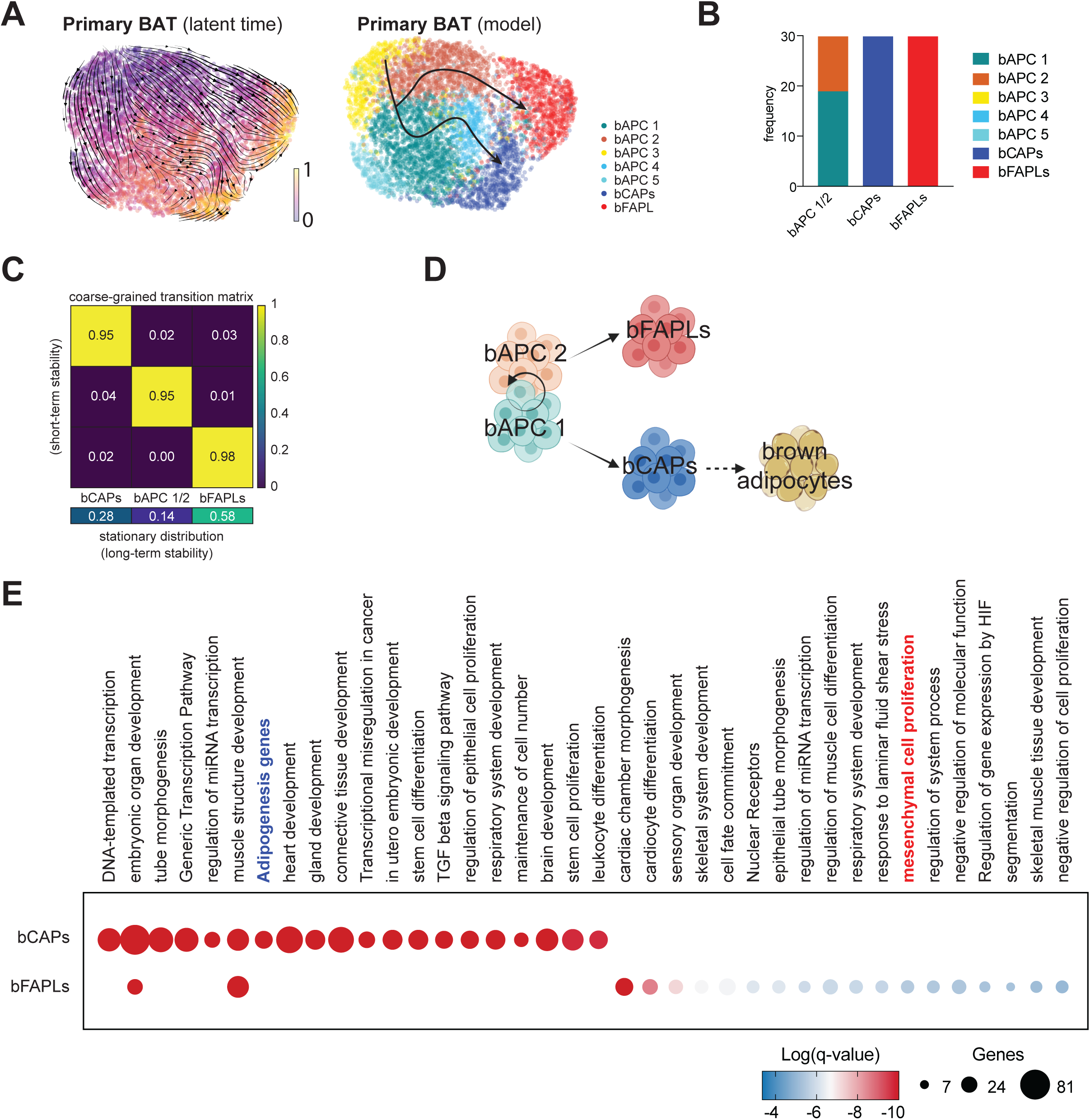
Pseudotime trajectory inferences and macrostate assignments in BAT. (**A**) Latent time UMAP visualization inferred by ScVelo and model of BAT APC differentiation trajectories highlight the transition from early precursors (bAPCs 1/2) to bCAPs and bFAPLs. (**B**) Composition of BAT APC macrostates (n = 30 cells) by CellRank2. (**C**) Coarse-grained transition matrix using CellRank2 GPCCA for WAT macrostates indicating the stability (diagonal elements) and transition probabilities (stationary distribution) amongst BAT macrostates. (**D**) Proposed developmental model of BAT APCs shows a mixed population of bAPC 1 and bAPC2 that can differentiate into bCAPs and bFAPLs, respectively. (**E**) Biological pathway analysis of the transcription factor drivers of bCAPs and bFAPLs shows enrichment in adipogenesis for bCAPs and mesenchymal-related pathways for bFAPLs.

**Figure EV5.**
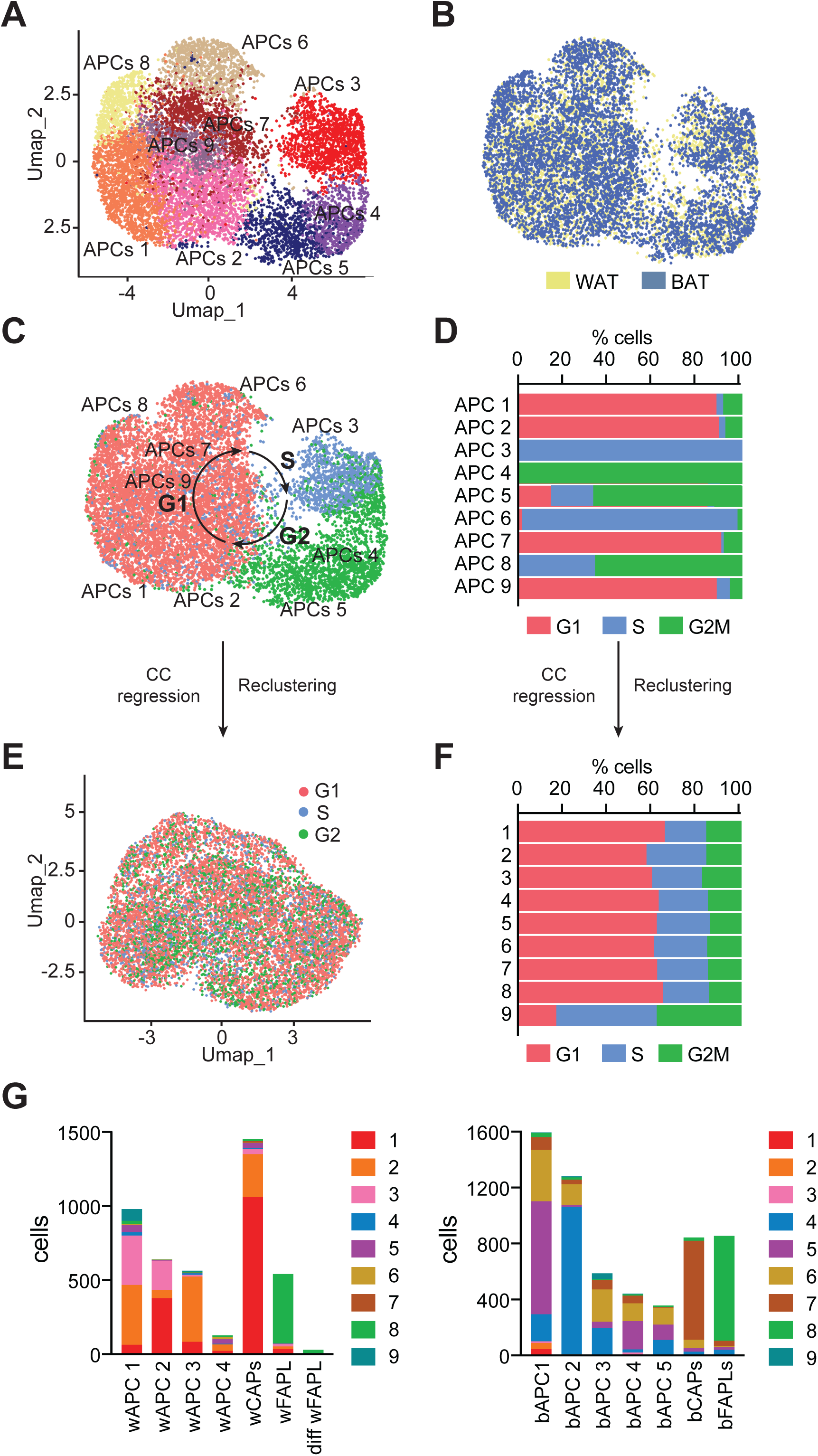
Integrated analysis of WAT and BAT precursor populations. (**A**) Clustering of integrated WAT and BAT APCs before cell cycle regression. (**B**) Distribution of WAT and BAT APCs in the aggregate UMAP. (**C**,**D**) Cell cycle phase (G1, S, G2/M) of WAT and BAT APCs and distribution across clusters. (**E**) UMAP of WAT and BAT APC aggregate after cell cycle regression. (**F**) Quantification of cell cycle frequency across clusters. (**G**) Cell count of white and brown APCs across each combined cluster.

**Figure EV6.**
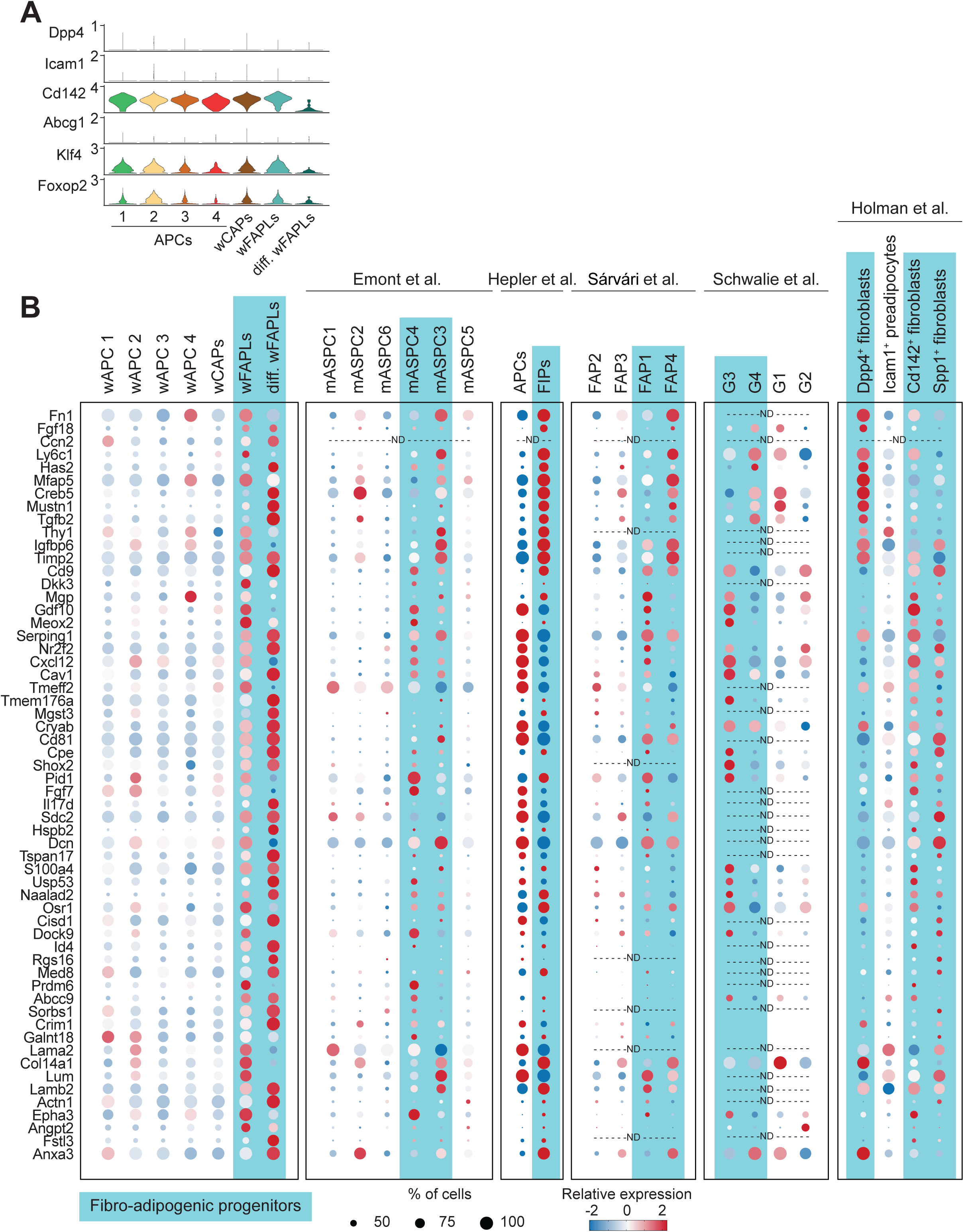
Identification of a shared FAP signature across multiple datasets. (**A**) Expression in culture WAT APCs of known genes previously reported to define mASPC4, FIP, Dpp4^+^, Icam^+^, Cd14^+^, Aregs and FAP populations in this WAT APC datasets. (**B**) Expression of other differentially expressed genes enriched in wFAPLs and differentiating wFAPLs compared to the expression found in previously reported fibro-adipogenic populations.

